# Drug repurposing: omeprazole increases the efficacy of acyclovir against herpes simplex virus type 1 and 2

**DOI:** 10.1101/313072

**Authors:** Martin Michaelis, Malte Christian Kleinschmidt, Mark N. Wass, Jindrich Cinatl

## Abstract

**Objectives:** Omeprazole was shown to improve the anti-cancer effect of the nucleoside-analogue 5-fluorouracil. Here, we investigated the effects of omeprazole on the activities of the antiviral nucleoside analogues ribavirin and acyclovir.

**Methods:** West Nile virus-infected Vero cells and influenza A H1N1-infected MDCK cells were treated with omeprazole and/ or ribavirin. Herpes simplex virus 1 (HSV-1)- or HSV-2-infected Vero or HaCat cells were treated with omeprazole and/ or acyclovir. Antiviral effects were determined by examination of cytopathogenic effects (CPE), immune staining, and virus yield assay. Cell viability was investigated by MTT assay.

**Results:** Omeprazole concentrations up to 80μg/mL did not affect the antiviral effects of ribavirin. In contrast, omeprazole increased the acyclovir-mediated effects on HSV-1- and HSV-2-induced CPE formation in a dose-dependent manner in Vero and HaCat cells. Addition of omeprazole 80μg/mL resulted in a 10.8-fold reduction of the acyclovir concentration that reduces CPE formation by 50% (IC_50_) in HSV-1-infected Vero cells and in a 47.7-fold acyclovir IC_50_ reduction in HSV-1-infected HaCat cells. In HSV-2-infected cells, omeprazole reduced the acyclovir IC_50_ by 7.3-fold (Vero cells) or by 12.9-fold (HaCat cells). Omeprazole also enhanced the acyclovir-mediated effects on viral antigen expression and virus replication in HSV-1- and HSV-2-infected cells. In HSV-1-infected HaCat cells, omeprazole 80μg/mL reduced the virus titre in the presence of acyclovir 1μg/mL by 1.6×10^5^-fold. In HSV-2-infected HaCat cells omeprazole 80μg/mL reduced the virus titre in the presence of acyclovir 2μg/mL by 9.2×10^3^-fold. The investigated drug concentrations did not affect cell viability, neither alone nor in combination.

**Conclusions:** Omeprazole increases the anti-HSV activity of acyclovir. As clinically well-established and tolerated drug, it is a candidate drug for antiviral therapies in combination with acyclovir.

## Introduction

Omeprazole and other proton pump inhibitors were found to increase the activity of anti-cancer drugs including the nucleoside analogue 5-fluorouracil (Luciani et al., 2004; Ikemura et al., 2017). Proton pump inhibitors are the most frequently prescribed drugs for the treatment and prophylaxis of gastroesophageal reflux as well as of gastric and duodenal ulcers that are associated with hyper-acidic states. Since they are known to be well-tolerated, they were suggested as repositioning candidates for the use as part of anti-cancer therapies (Ikemura et al., 2017).

Here, we investigated the effects of omeprazole on the efficacy of the antiviral nucleoside analogues acyclovir and ribavirin. We found that omeprazole enhanced the antiviral effects of acyclovir against herpes simplex virus type 1 (HSV-1) and HSV-2 but did not influence the activity of ribavirin against West-Nile viruses or influenza viruses.

## Materials and methods

### Cell culture

Vero and MDCK cells were obtained from the American Type Culture Collection (ATCC, Rockville, MD) and cultured at 37° C in minimum essential medium (MEM) supplemented with 10% foetal bovine serum. HaCaT cells were purchased from CLS Cell Line Services GmbH (Eppelheim, Germany) and cultivated in Iscove’s modified Dulbecco’s medium (IMDM) supplemented with 10% foetal bovine serum.

### Viruses

HSV-1 strain McIntyre and HSV-2 strain MS were both obtained from ATCC. West Nile virus (WNV) strain NY385-99 was kindly provided by Dr J ter Meulen (Institut für Virologie, Phillips-Universität, Marburg, Germany). Virus stocks were prepared in Vero cells grown in MEM with 4% foetal bovine serum. The influenza virus strain Influenza A/New Caledonia/20/99 (H1N1) was received from the WHO Influenza Centre (National Institute for Medical Research, London, UK). Virus stocks were prepared in MDCK cells grown in 4% foetal bovine serum. Infectious virus titres were determined by titration on MDCK cell monolayer in 96-well plates as 50% tissue culture infectious dose (TCID_50_) by the method of Spearman and Kärber (Spearman, 1908; Kärber, 1931).

### Drugs

Acyclovir was received from GlaxoSmithKline (Munich, Germany), omeprazole from AstraZeneca (Wedel, Germany), and ribavirin from Valeant Pharmaceuticals Germany GmbH (Eschborn, Germany).

### Cytopathogenic effect (CPE) reduction assay

For the investigation of HSV-1- and HSV-2-induced cytopathogenic effects (CPEs), confluent Vero or HaCaT cell monolayer in 96-well microtiter plates were inoculated with HSV-1 or HSV-2 at MOI 1 or 0.1, respectively. Following a 1h incubation period, the inoculum was removed and the drugs, either alone or in combination, were added. The virus-induced CPE was recorded microscopically after 48h post infection.

For the investigation of WNV-induced CPEs, Vero cell monolayers were infected with MOI 0.1. Following a 1h virus incubation period, the medium was removed and infected cells were incubated in medium containing different concentrations of drugs at the respective concentration. The CPE was recorded at 48h post infection.

Confluent MDCK cell monolayers were infected with Influenza H1N1 (MOI 0.01). Following a 1h virus incubation period, the medium was removed and infected cells were incubated in medium containing different concentrations of drugs at the respective concentration. The CPE was recorded at 24h post infection. CPEs were scored by two independent examiners and expressed in % of the untreated virus control that was defined to be 100%.

### Immunostaining

Intracellular HSV protein was evaluated by immunostaining. Cells were fixed with 60/40 ice cold methanol/acetone for 15 min. Staining was performed using a rabbit polyclonal antibody directed against HSV-1 (ab9533) and a sheep polyclonal antibody directed against HSV-2 (ab21112) in combination with biotin-conjugated secondary goat anti-rabbit (ab6720) and rabbit anti-sheep (ab6746) antibodies (all antibodies derived from Abcam, Cambridge, UK). Protein was visualised using streptavidin-peroxidase complex with AEC as a substrate.

### Viability assay

The cellular viability was assessed on confluent cell layers with the 3-(4,5-dimethyl-2-thiazolyl)-2,5-diphenyl-2H-tetrazolium bromide (MTT) assay method as described previously (Michaelis et al., 2007). The viability was expressed as percentage of non-treated control.

## Results

### Effects of omeprazole in combination with ribavirin on cytopathogenic effect (CPE) formation in WNV- or influenza A H1N1-infected cells

Omeprazole 80μg/mL did not affect the effects of ribavirin on CPE formation in WNV-infected Vero cells or H1N1-infected MDCK cells (Figure 1A, Suppl. Table 1).

**Figure 1.**
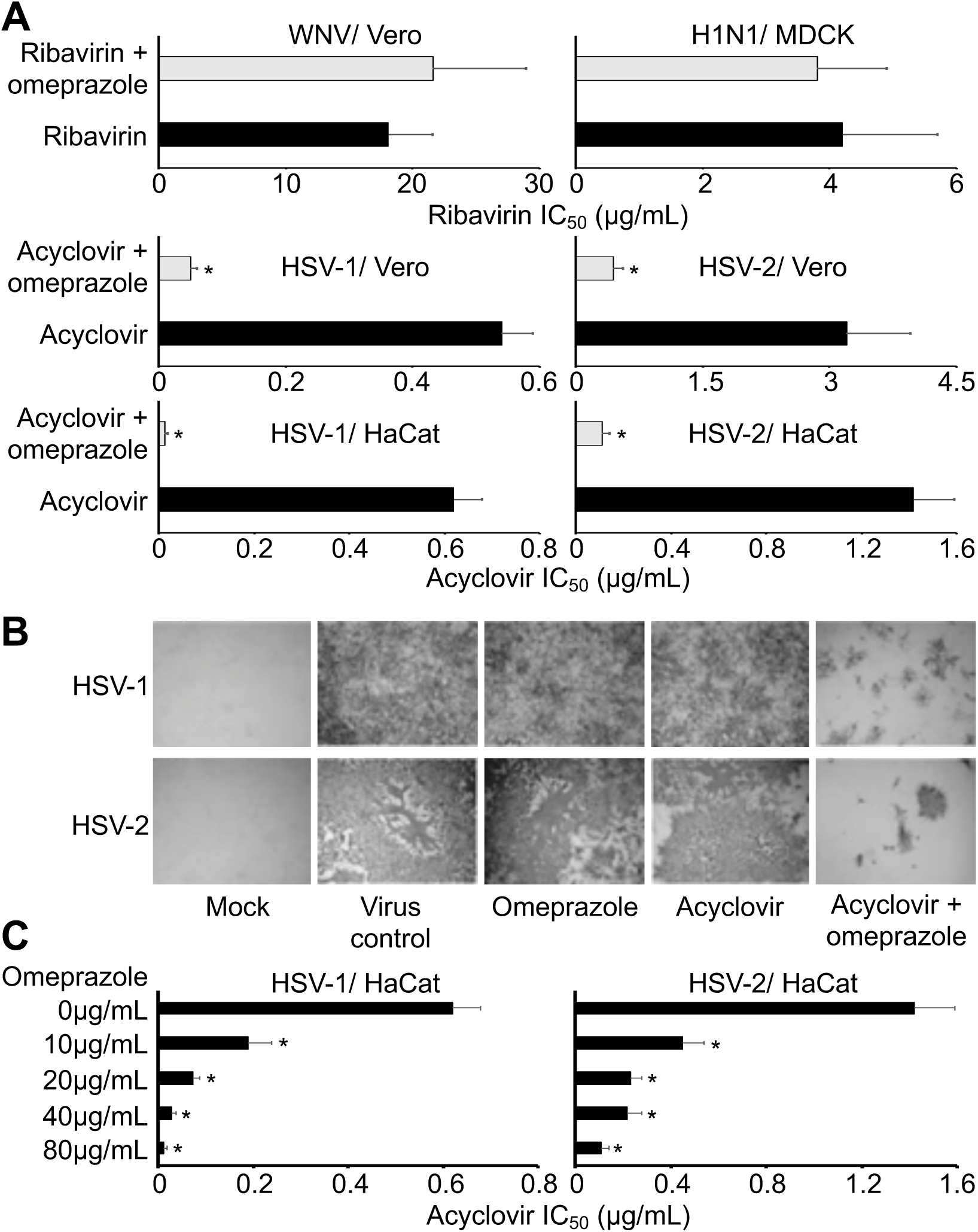
Cytopathogenic effect (CPE) formation and viral gene expression in the presence of omeprazole antiviral nucleoside analogues. A) Effects of omeprazole (80μg/mL) on the concentrations of antiviral nucleoside analogues that reduce CPE formation by 50% (IC_50_) using West Nile virus (WNV)-infected Vero cells, influenza A H1N1-infected MDCK cells, and HSV-1- or HSV-2-infected Vero or HaCat cells. Omeprazole alone did not reduce CPE formation. Numerical values are presented in Suppl. Table 1. B) Effects of omeprazole and acyclovir on the expression of virus proteins in HSV-1- and HSV-2-infected Vero cells. HSV-1-infected cells were treated with omeprazole 80μg/mL and/ or acyclovir 0.31μg/mL. HSV-2-infected cells were treated with omeprazole 40μg/mL and/ or acyclovir 0.6μg/mL. C) Concentration-dependent effects of omeprazole on the acyclovir IC_50_ in HSV-1- or HSV-2-infected HaCat cells as determined by CPE formation. Numerical values are presented in Suppl. Table 2. The investigated drug concentrations did not affect cell viability, neither alone nor in combination. ^*^ P < 0.05 relative to nucleoside analogue alone

### Effects of omeprazole in combination with acyclovir on HSV-1 and HSV-2 replication

In the presence of omeprazole 80μg/mL, acyclovir concentrations that reduced CPE formation by 50% (IC_50_) were reduced by 11-fold in HSV-1-infected Vero cells and by 7-fold in HSV-2-infected Vero cells. In addition, omeprazole 80μg/mL reduced the acyclovir IC_50_s by 48-fold in HSV-1-infected HaCat cells and by 13-fold in HSV-2-infected HaCat cells (Figure 1A, Suppl. Table 1). Immune staining also indicated reduced numbers of virus-infected cells after treatment with a combination of omeprazole and acyclovir compared to either single treatment (Figure 1B). Further experiments indicated that omeprazole reduced acyclovir IC_50_s in HSV-1- and HSV-2-infected HaCat cells in a dose-dependent fashion (Figure 1C, Suppl. Table 2).

The detection of virus titres revealed moderate effects of omeprazole on virus replication. Again, omeprazole strongly increased the anti-HSV-1 and anti-HSV-2 effects of acyclovir (Figure 2, Suppl. Table 3). The investigated omeprazole and acyclovir concentrations did not affect cell viability, neither alone nor in combination.

**Figure 2.**
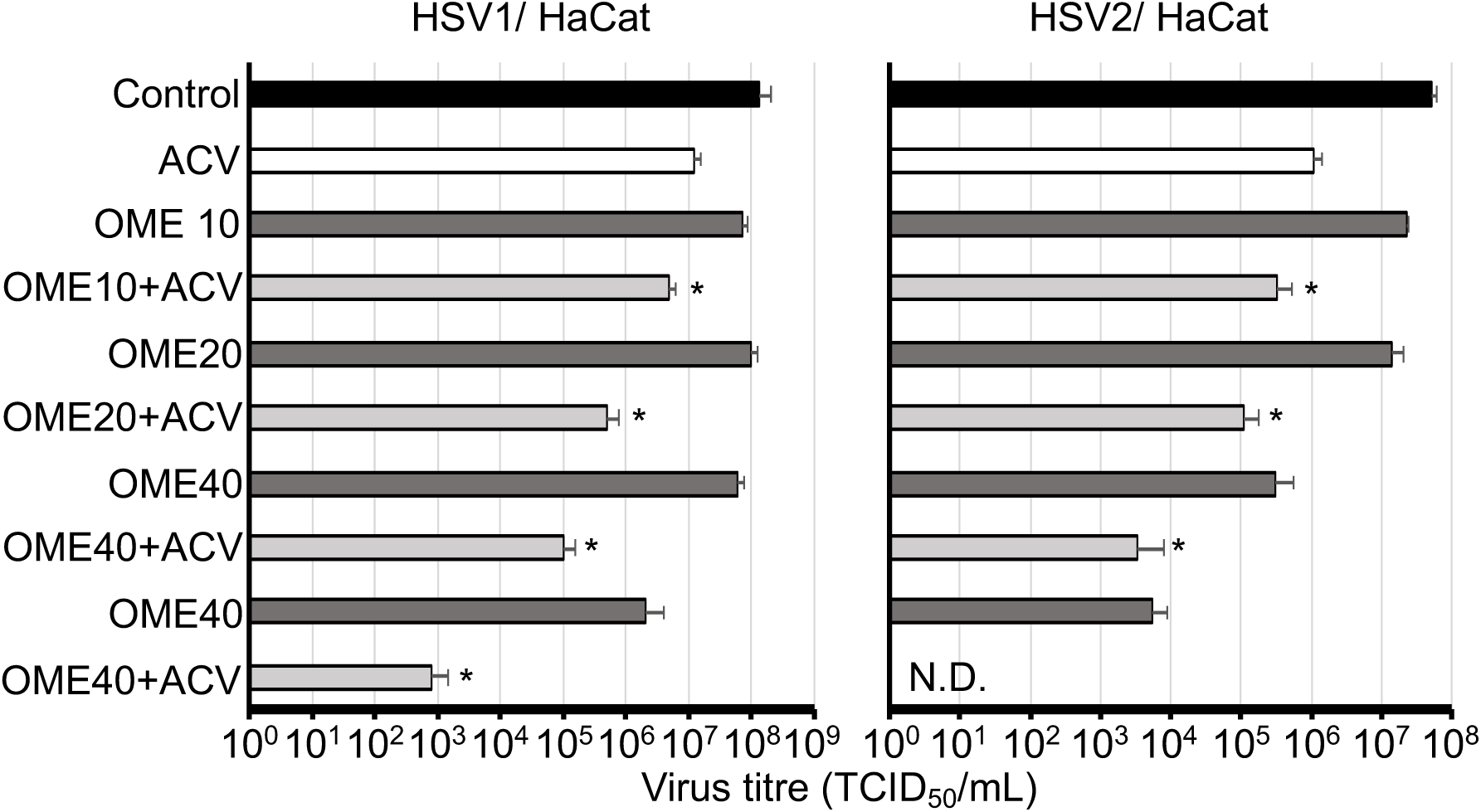
Effect of acyclovir 1μg/mL (HSV-1) or 2μg/mL (HSV-2) alone or in combination with varying omeprazole (OME) concentrations (μg/mL) on HSV-1 and HSV-2 titres in HaCat cells. Numerical values are presented in Suppl. Table 3. ^*^ P < 0.05 relative to acyclovir alone; N.D. = no detectable virus titre

## Discussion

Based on previous investigations that showed that omeprazole increases the anti-cancer activity of the nucleoside analogue 5-fluorouracil (Luciani et al., 2004), we here investigated the effects of omeprazole on the antiviral effects of ribavirin and acyclovir. Ribavirin is a broad spectrum antiviral drug (Beaucourt & Vignuzzi, 2014). However, omeprazole did neither modify ribavirin-mediated effects in H1N1 influenza A virus-infected cell cultures nor in West Nile virus-infected cell cultures.

Acyclovir is a first line drug for HSV-1, HSV-2, and varicella zoster virus infection (Piret & Boivin, 2016; Klysik et al., 2018). In contrast to the lack of effect of omeprazole on ribavirin-mediated antiviral effects, omeprazole interfered with HSV-1 and HSV-2 replication in a dose-dependent fashion. Omeprazole increased the anti-HSV activity of acyclovir in Vero cells and in the human keratinocyte cell line HaCat. Since omeprazole and acyclovir were both added simultaneously to virus-infected cell cultures after a 1h viral adsorption period, omeprazole increases the effects of acyclovir during the viral replication cycle.

The mechanism by which omeprazole enhances the activity of acyclovir seems to differ from the mechanism by which the compound increases 5-fluorouracil efficacy. Omeprazole pre-treatment was necessary to increase 5-fluorouracil activity (Luciani et al., 2004). In contrast, the combination of omeprazole and acyclovir exerted its combined activity when added at the same time.

There is a need for improved therapies for HSV-1- and HSV-2-associated disease. After primary infection, HSV-1 and HSV-2 establish life-long persistence which may result in recurrent disease which typically manifests as herpes labialis or herpes genitalis and which may be associated with significant morbidity (Gnann & Whitley, 2016; Heslop et al., 2016; Klysik et al., 2018). Even in the case of herpes labialis, which is not commonly associated with complications, treatment success is not always satisfactory as indicated by the introduction of topical acyclovir/ hydrocortisone combinations (Nguyen et al., 2014). In immunodeficient individuals, HSV-1/-2 infection is often associated with more severe disease, and resistance formation to acyclovir is a severe problem (Piret & Boivin, 2016; Karrasch et al., 2018). Moreover, ocular HSV infection is a major cause of blindness in industrialised countries (Klysik et al., 2018). In conclusion, omeprazole substantially enhances the antiviral effects of acyclovir in HSV-1- and HSV-2-infected cells. Improved therapies for HSV-1/2 infection are highly desirable, in particular for immunocompromised individuals. Since omeprazole is a clinically well-established drug with a preferable safety profile, it is an excellent candidate for drug repositioning strategies (Ikemura et al., 2017).

